# Trophic level associated gut length divergence evolved under sexual conflict in Lake Malawi cichlids

**DOI:** 10.1101/2024.12.09.627601

**Authors:** Aldo Carmona Baez, Patrick J. Ciccotto, Emily C. Moore, Erin N. Peterson, Melissa S. Lamm, Natalie B. Roberts, Kaitlin P. Coyle, M. Kaitlyn Barker, Ethan Dickson, Amanda N. Cass, Guilherme S. Pereira, Zhao-Bang Zeng, Rafael F. Guerrero, Reade B. Roberts

## Abstract

Variation in gastrointestinal morphology is associated with dietary specialization across the animal kingdom. Gut length generally correlates with trophic level, and increased gut length in herbivores is a classic example of adaptation to cope with diets with lower nutrient content and a higher proportion of refractory material. However, the genetic basis of gut length variation remains largely unstudied, partly due to the inaccessibility and plasticity of the gut tissue, as well as the lack of dietary diversity within traditional model organisms relative to that observed among species belonging to different trophic levels. Here, we confirm the genetic basis of gut length variation among recently evolved Lake Malawi cichlid fish species with different dietary adaptations. We then produce interspecific, inter-trophic-level hybrids to map evolved differences in intestinal length in an F2 mapping cross between *Metriaclima mbenjii*, an omnivore with a relatively long gut, and *Aulonocara koningsi*, a carnivore with a relatively short gut. We identify numerous candidate quantitative trait loci for evolved differences in intestinal length. These quantitative trait loci are predominantly sex-specific, supporting an evolutionary history of sexual conflicts for the gut. We also identify epistatic interactions potentially associated with canalization and the maintenance of cryptic variation in the cichlid adaptive radiation. Overall, our results suggest a complex, polygenic evolution of gut length variation associated with trophic level differences among cichlids, as well as conflicts and interactions that may be involved in evolutionary processes underlying other traits in cichlids.

**Summary:** This study examines the genetic basis of gut length variation in Lake Malawi cichlids, which exhibit different dietary adaptations. It highlights how cichlids recapitulate a broad taxonomic trend: gut length correlates with trophic level, with herbivores and omnivores having longer intestines than carnivores. By creating hybrids of *Metriaclima mbenjii* (omnivore) and *Aulonocara koningsi* (carnivore), we identify several quantitative trait loci and epistatic interactions underlying gut length differences. These genetic associations are predominantly sex-specific, suggesting historical sexual conflicts. The results indicate complex, polygenic evolution of gut morphology in these fish, and suggest evolutionary interactions and processes shaping dietary traits across species.

## Introduction

Adaptation to different diets is a major driver of species and phenotypic diversity, and requires the concomitant evolution of a suite of traits, including morphology, sensory systems, behavior, metabolism, and the digestive system. Phenotypic variation in the gastrointestinal system is generally predictable across taxonomic groups, where the length and morphological complexity of the gut scale with trophic level and the processing requirements for particular diets (Duque-Correa et al., 2021; German and Horn 2006; Griffen and Mosblack, 2011). Generally, carnivores eating an easily digestible, nutrient-rich diet have shorter and simpler guts than herbivores consuming relatively nutrient-poor diets or those requiring the processing of refractory material (Karasov and Douglas, 2013; Starck, 2005). Despite our extensive knowledge of the variability of the morphology and physiology of the gastrointestinal system across vertebrate taxa, the genetic components underlying this naturally evolved phenotypic diversity remain unknown.

Bridging this knowledge gap, however, poses challenges. First, traditional genetic model systems such as mouse (*Mus musculus*) and zebrafish (*Danio rerio*) lack the diet-related phenotypic diversity observed in the gastrointestinal tract among vertebrate species in the wild. Second, sampling the gut requires invasive procedures, which force morphological and physiological studies to use postmortem samples that are challenging to obtain (Greene and McKenney et al., 2022; Yawitz et al., 2022). Third, the gastrointestinal tract displays high phenotypic plasticity, particularly in response to dietary differences and other environmental factors that trigger changes in development or homeostasis, which in turn impact the morphology and physiology of adult organisms (Olsson et al., 2007; Naya et al., 2007; Pfennig, 1992). Because of these reasons, appropriate model systems for the study of the genetic basis of gastrointestinal dietary adaptations would ideally exhibit: a) phenotypic diversity in the gastrointestinal tract stemming from dietary adaptation; b) close relatedness among the species/populations composing the model system in order to be able to perform forward genetic experiments via comparative genomics or quantitative trait loci (QTL) mapping; c) limited body size divergence among populations to avoid allometry-related issues given the correlation between gut and body size; and d) laboratory tractability, so experimental organisms can be raised and sampled under controlled conditions in order to reduce the confounding effects of phenotypic plasticity, particularly for diet. Examples of established natural model systems that appear to meet these criteria are deer mice (*Peromyscus*), the Mexican tetra (*Astyanax mexicanus*), and cichlid fish (Yawitz et al., 2022; Riddle et al., 2021; Feller et al., 2022).

In this work, we study the genetic basis of gut length in the context of the adaptive radiation of Lake Malawi cichlid fishes. This radiation consists of more than 800 recently diverged (∼1M years ago) species that have specialized to a broad range of food sources, reflected in the abundant phenotypic diversity observed in behavior, craniofacial morphology, and sensory traits (Brawand et al., 2014). Additionally, previous studies have shown that Malawi cichlids sampled in the wild display differences in gut length associated with trophic levels, recapitulating trends observed in other, broader taxonomic comparisons (Fryer and Iles, 1972; Reinthal, 1989). Furthermore, the parallel Lake Tanganyika cichlid radiation (phylogenetically proximate to Malawi cichlids) also shows differences in gut length associated with trophic level (Wagner et al., 2009). However, comparative gut length data for cichlid species raised under common diet and housing conditions are limited, making it difficult to disentangle genetic versus environmental impacts on the trait.

Alongside adaptation to dietary differences, other pressures such as habitat adaptation, sexual selection, and sexual conflicts (or combinations of these processes) have played key roles in the evolutionary history of African cichlids and have likely impacted gut evolution.

Indeed, recent studies comparing traits within single cichlid species identified sex-specific differences in two gastrointestinal traits: gut microbiota composition and intestinal length (Faber- Hammond et al., 2019; Moore et al., 2022). This sex-associated trait variation may be linked to sex differences in life history, including sexually dimorphic territoriality and pronounced differences in parental investment (Ribbink, 1983). Particularly notable in Lake Malawi cichlids is their use of maternal mouthbrooding, where eggs and larval fish are incubated in their mother’s mouth until they develop into free-swimming fry. Maternal mouthbrooding results in repeated instances of prolonged self-induced starvation limited to a single sex, thus creating sex differences in dietary intake that may require sex differences in gut biology to maintain fitness (Konings, 2007; Faber-Hammond et al., 2019; Moore et al., 2022).

Here, we first demonstrate that trophic-level gut length variation in Malawi cichlid fishes has a genetic component by raising a multi-species panel under controlled laboratory conditions. We then perform QTL mapping in an F2 population from a hybrid cross between two sympatric cichlid species, the omnivorous *Metriaclima mbenjii,* a generalist whose diet includes algae, plankton, and invertebrates, and the carnivorous species *Aulonocara koningsi,* a specialized invertivore. This study complements previous work using similar strategies to map trophic traits in Malawi cichlids, such as jaw morphology and vision (Albertson et al., 2003; O’Quin et al., 2010). To our knowledge, ours is one of only a few recent studies characterizing the genetic architecture of gut length in a non-agricultural and wild-derived model system (for others, see Riddle et al., 2021, and Feller et al., 2022). As such, these findings are foundational for understanding basic gut biology and the genetic evolution of the gastrointestinal tract in vertebrate species.

## Methods

### Animal husbandry

All animal research was conducted under protocol 14-101-O, as approved by the IACUC at North Carolina State University. All individuals (including hybrids) used for gut length measurements were bred and raised in a common recirculating system under the same laboratory conditions. All fish were fed a 1:1 combination of Premium Super Brine Shrimp Flake and Premium Vegetable Flake (Ken’s Fish and Pet Supplies; Taunton, MA), consisting of 42.5% protein, 9.5% fat, 5% fiber, and 9% moisture. All individuals were collected as eggs, hatched in 3 L tanks, and fed ∼25 mg of flake mix twice a day starting at 21 dpf. At 45 dpf, they were transferred to 10 L tanks and fed ∼100 mg of flake mix twice a day until 90 dpf, when they were transferred to 37.9 L tanks and fed ∼200 mg of flake mix twice a day. Throughout, fish were housed at 5 to 12 individuals per tank. Individuals were sampled at 22 weeks of age following two days of fasting. A hybrid cross was produced from a single *Metriaclima* female crossed to two *Aulonocara* males; the inclusion of the second grandsire was inadvertent and resulted from an unexpected fertilization event in these species with external fertilization. The resulting F1 hybrids were intercrossed to produce an F2 mapping population (*N* = 495).

### Dissections

All fish were euthanized in a solution of 250 mg/L tricaine methanesulfonate (Tricane-S, Western Chemical, Ferndale, WA, U.S.A.), prior to dissection for removal of the alimentary canal. Intestines were unwound and laid in a straight line on a flat surface in a consistent manner, with care not to stretch the tissue and without the use of relaxants to eliminate contraction activities. Gut length (GL) was measured as intestinal length from the posterior of the stomach to the anus, and standard length (SL) was measured from the tip of the snout to the posterior margin of the hypural plate using Vernier calipers. Phenotypic sex was determined at dissection by removing and squashing a portion of the gonad on a slide to confirm the presence of oocytes or mature sperm under a microscope. Sex data at dissection was further compared to genetic marker information (see below).

### Data analysis of multi-species panel

Species were scored as carnivorous, omnivorous, or herbivorous based on information from Konings (2007). Species that primarily feed on fish and/or macroinvertebrates were scored as carnivorous, species that feed on some combination of macroinvertebrates, algae, and plankton were scored as omnivorous, and species that primarily feed on algae and/or macrophytes were scored as herbivorous. While there is further variation in diet specializations within these trophic groupings, our survey did not include enough species for finer comparisons. See Supplementary Table 1 for the list of species and numbers analyzed.

In order to test for differences in gut length across trophic levels while accounting for allometry, we performed a one-way ANOVA test on the residuals of the model *GL∼SL* followed by a post hoc Tukey-Kramer HSD test. This analysis was performed in the JMP Pro 17 (SAS) statistical platform. Visualizations of the multi-species panel were generated with the R statistical platform (version 4.0.3; R Core Team, 2021) and the packages ggplot2, tidyr, and dplyr (Wickham et al., 2019).

### Genetic map building

Libraries for double-digest restriction-site associated DNA sequencing (ddRADseq) were produced and processed as previously reported in DeLorenzo et al., 2023. The genetic map (Supplementary Fig. 1) was built on the R statistical platform (version 4.0.3; R Core Team, 2021) with the package R/QTL (Broman et al, 2003). The RAD markers were sorted and binned in linkage groups according to their position in the *M. zebra* UMD2a reference genome, using corresponding chromosome numbering (Conte et al., 2019). Markers with more than 20% of missing genotypes on autosomes and markers with more than 40% of missing genotypes on unplaced scaffolds were removed from the dataset. A chi-square test was performed with the function geno.table() to detect markers with distorted segregation patterns; markers with a Bonferroni-corrected p-value < 0.05 were removed. A subset of markers on linkage group 11 showed segregation distortion, in a pattern consistent with a potential recessive lethal allele in the families descended from one F1 dam. Based on this observation, the chromosome 11 genotypes from the individuals displaying segregation distortion were masked by setting them as missing data. After masking, the genetic markers on chromosome 11 matched the expected 1:2:1 segregation pattern (Supplementary Table 2). The pairwise recombination frequencies among markers was calculated and an initial map was estimated with the functions est.rf() and est.map(), respectively. Markers whose recombination frequency profile did not match their position in the genetic map, likely due to being located in structural variants or misassembled regions of the reference genome, were relocated so that the number of crossovers was minimal. Individuals with more than 350 crossovers (4 out of 495) were removed from the dataset.

Markers that increased the size of the map by 6 cM or more and were within 3 Mb from their flanking markers were considered to be inflating the map and were excluded. Markers located in unplaced scaffolds were integrated into a given linkage group if they had recombination frequency values < 0.15 with at least 5 markers from that linkage group. The function calc.errorlod() was used to detect genotyping errors and genotypes with a LOD score >= 3 were set as missing data. The map was pruned using a non-overlapping window algorithm that selected the marker in a given 2cM window with the least missing data. The remaining markers were manually curated and the missing genotypes were set as not-AA or not-BB if there were at least 3 sequencing reads supporting a particular allele. The final map was estimated and the maximum likelihood estimate of the genotyping error rate (0.0001) was obtained with the function est.map(). The resulting map corresponds to 22 chromosomes and uses conventional numbering for the standard Lake Malawi cichlid karyotype (Conte et al., 2019).

### Data analysis of phenotypic data in hybrid cross

As mentioned above, the sex of the F2 individuals was first obtained through gonad dissection data. Of 491 F2 individuals that passed genotype filters, 145 had ambiguous sex calls based on dissection notes. Sex in the F2 cross is controlled by a previously-described XY sex determination locus on chromosome 7 (Ser et al., 2010), which we confirmed as the sole sex determination locus through QTL mapping (Fig. 2a). In order to infer the sex of individuals with ambiguous dissection data, the dissection information was complemented with information from two genetic markers in close proximity to the mapped XY sex determination locus (see also QTL mapping methods). The sex-linked markers were located on chromosome 7, at 49,393,905 bp and 52,246,354 bp in the *M. zebra* UMD2a reference genome (Conte et al., 2019). An F2 individual was classified as female only if the dissection notes mentioned the presence of ovaries or eggs in the gonad and the marker data suggested the absence of the Y-linked allele. Similarly, an individual was classified as male if the dissection notes mentioned the presence of testis or sperm in the gonad and the marker data suggested the presence of the Y-linked allele. The final number of individuals with high-confidence sex calls was 448 (218 females and 230 males), displaying a 1:1.03 female-to-male ratio. The remaining 43 individuals with ambiguous sex calls were excluded from subsequent analyses.

Exploratory data analysis of sex, standard length and gut length was performed using the R statistical platform (version 4.0.3; R Core Team, 2021) and the packages ggplot2, tidyr, and dplyr (Wickham et al., 2019).

### QTL mapping analyses

QTL mapping analyses were performed using Windows QTL Cartographer version 2.5 (Silva et al., 2012). Sex was mapped using an Interval Mapping (IM) approach using the data of 346 individuals that had unambiguous sex calls at dissection. Other mapping analyses used a set of 448 individuals with sex calls determined as described above.

In order to map gut length while accounting for allometry, the residuals of the model *GL∼SL* were mapped using an IM approach. Additionally, GL and SL were mapped using a multi-trait IM approach (Jiang and Zeng, 1995) that consisted of mapping SL, GL, and the joint trait SL-GL. The QTL mapping analyses of the residuals, SL, GL, and the joint trait SL-GL were performed on three different subsets of the data: all individuals, only female individuals, and only male individuals. The analyses performed on the "all individuals" subset included sex as a covariate (Otrait in Windows QTL Cartographer). Similarly, chromosome 7, where the XY sex determination system is located (Ser et al., 2010), was removed from the analyses performed on the female and male subsets since splitting the data by sex produces segregation distortion on this chromosome. The significance threshold for the single-trait scans (SL, GL, residuals) was set to LOD=3, while the significance threshold for the joint trait SL-GL was set to LOD=4. Allele effects of candidate QTL were based on the marker position at each QTL that was most frequently identified in each chromosome across the different IM scans (Supplementary Table 3).

In order to further refine the location, number, and effect sizes of the QTL underlying our traits of interest, we performed Multiple Interval Mapping (MIM) analyses of SL, GL, and the residuals for the model *GL∼SL* using Windows QTL Cartographer version 2.5 (Silva et al., 2012). The analysis was performed on three different subsets of the data: all individuals, only female individuals, and only male individuals. In contrast to the IM analyses, the joint trait SL-GL was not mapped since multi-trait MIM analysis has not been implemented for F2 mapping populations on Windows QTL Cartographer. In addition, the analyses performed on the "all individuals" subset did not include sex as a covariate since this option is not implemented in the MIM procedure of Windows QTL Cartographer. Chromosome 7 was removed in the analyses performed on the female and male subsets, as in the IM analyses.

The MIM analyses consisted of the following steps: 0) The 9 candidate QTL selected from the IM procedure were used as the initial model. 1) The significance of the main QTL was tested and the non-significant QTL were removed from the model. In the case of the first iteration, if none of the initial model QTL were maintained, a scan to look for other QTL was performed. The significance of QTL interaction terms was also assessed if epistatic terms were present in the model. QTL interaction terms were removed if they did not reach significance. 2) The positions of the QTL were optimized. 3) A scan was performed for new main QTL and QTL interactions (between QTL already present in the model). 4) New QTL were added to the model by repeating steps 1-3. The search for new QTL stopped when the model had seven or more main effect QTL at the start of a new iteration or when the model reached convergence. The statistical criterion used in the model selection procedure was Bayesian information criterion "Model 0" (BIC-M0; see definition below), a relatively strict criterion. Additionally, the residuals and SL MIM analyses in the male and female subsets were replicated using a more lenient approach by alternating the use of two criteria: BIC-M0 and Akaike information criterion (AIC). Information criterion (IC) values are given by 𝐼𝐶 = 𝑛 x 𝑙𝑛(𝑄 x 𝑄) + 𝑝 x 𝑐(𝑛), where 𝑛 represents the sample size, 𝑙𝑛(𝑄 x 𝑄) the residual variance of the model, 𝑝 the number of regressors (markers) in the model, and 𝑐(𝑛) a factor that determines the stringency of the penalty for model complexity. The BIC-M0 sets 𝑐(𝑛) = 𝑙𝑛(𝑛), and AIC sets 𝑐(𝑛) = 2 (Windows QTL Cartographer version 2.5, Silva et al., 2012). For all MIM analyses, the walking speed was 1 cM, and the window size 10 cM.

The location, effect, and significance of the identified QTL were recorded in tables to allow for comparison across analyses. QTL identified on different scans that were less than 10 cM apart and had similar magnitudes and directions were reported in a single row in the summary tables. QTL coordinates can be found in the Supplementary Tables 3, 4, and 5. The QTL LOD score profiles and QTL effects, as well as two epistatic interactions, were visualized with ggplot2 (Wickham et al., 2019), R/QTL (Broman et al., 2003), and in-house R scripts (Fig. 4, Supplementary Figs. 2, 3, and 4).

## Results

### Phenotypic analysis

We bred and raised thirteen cichlid species in the lab, representing different dietary specializations and broadly classified as carnivores, omnivores, or herbivores (see Methods). All were provided with the same diet and housing densities to minimize environmental effects on variance in gut length. Under these common laboratory conditions, the species recapitulate trends reported in natural populations, with carnivores having significantly shorter intestines than omnivores and herbivores when grouped by trophic level and controlling for body size by using residuals of the model *GL∼SL* (Fig. 1a and b, Supplementary Table 1; one-way ANOVA, F = 60.8, p < 0.0001); omnivore and herbivore intestine lengths are not significantly different. We note that one genus in our survey, *Labidochromis*, includes two species adapted to two different trophic levels, and their gut length diverges as expected based on trophic level (*t-*test, F = 7.33, p = 0.014). When all species are collapsed by trophic level classification, intestine length is not significantly different between omnivore and herbivore classifications, but each significantly differs from carnivores (Tukey’s HSD test; Fig. 1b).

**Fig. 1.**
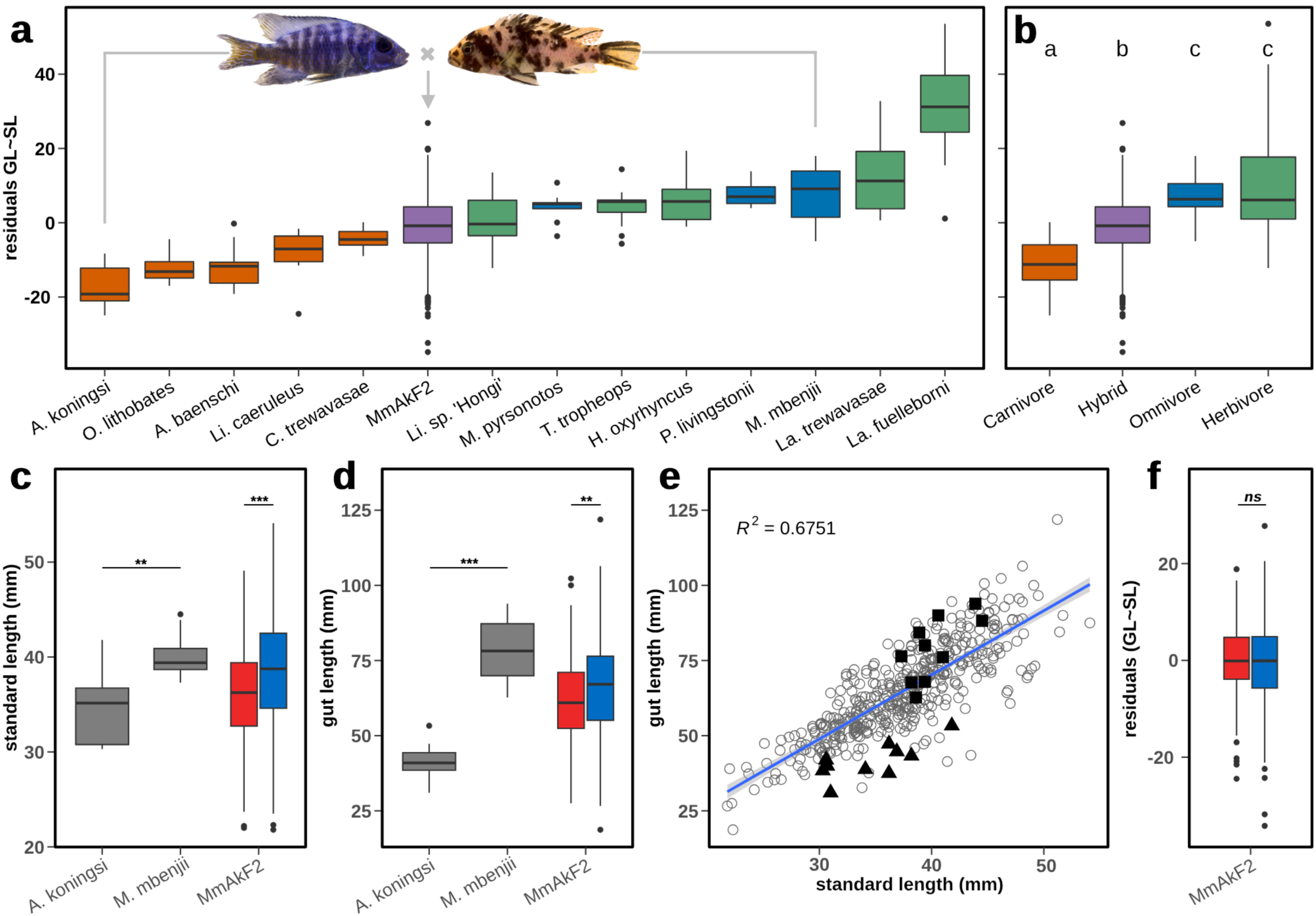
Phenotypic analysis of body standard length and gut length data. a) Phenotypic survey of individuals raised under common diet and housing conditions supports a genetic basis of normalized intestine length variation by species; see Supplementary Table 1 for numbers tested and statistical comparisons. Inset fish images are representative individuals of parental species for hybrid cross. b) Binning intestine length data by trophic level of species shows that carnivores (orange) have significantly shorter intestines than omnivores (blue) or herbivores (green), but omnivores and herbivores are not significantly different from each other. The hybrid F2 mapping population (MmAkF2, purple) exhibits intermediate normalized intestine length distinct from both carnivore and omnivore-herbivore species bins (ANOVA with Tukey’s HSD, different letters indicate significance at p < 0.05). c) Standard body length and d) gut length in the parental species (N = 10 each) of the hybrid cross, and female (red, N = 218) and male (blue, N = 230) individuals of the hybrid F2 population (*t*-test, *** p < 0.001, ** p < 0.01). e) Scatter plot of standard length versus gut length for individuals in the F2 mapping population (circles), and the parental samples (squares: *M. mbenjii*; triangles: *A. koningsi*). The linear regression line of the model GL∼SL with confidence interval is shown for the F2 individuals. f) Residuals of the model GL∼SL are not significantly different by sex in the F2 hybrids. All sampled F2 individuals (N = 495) are presented in a and b. Only the F2 individuals with unambiguous sex calls and used for mapping (N = 448) are included in c-e.

**Fig. 2.**
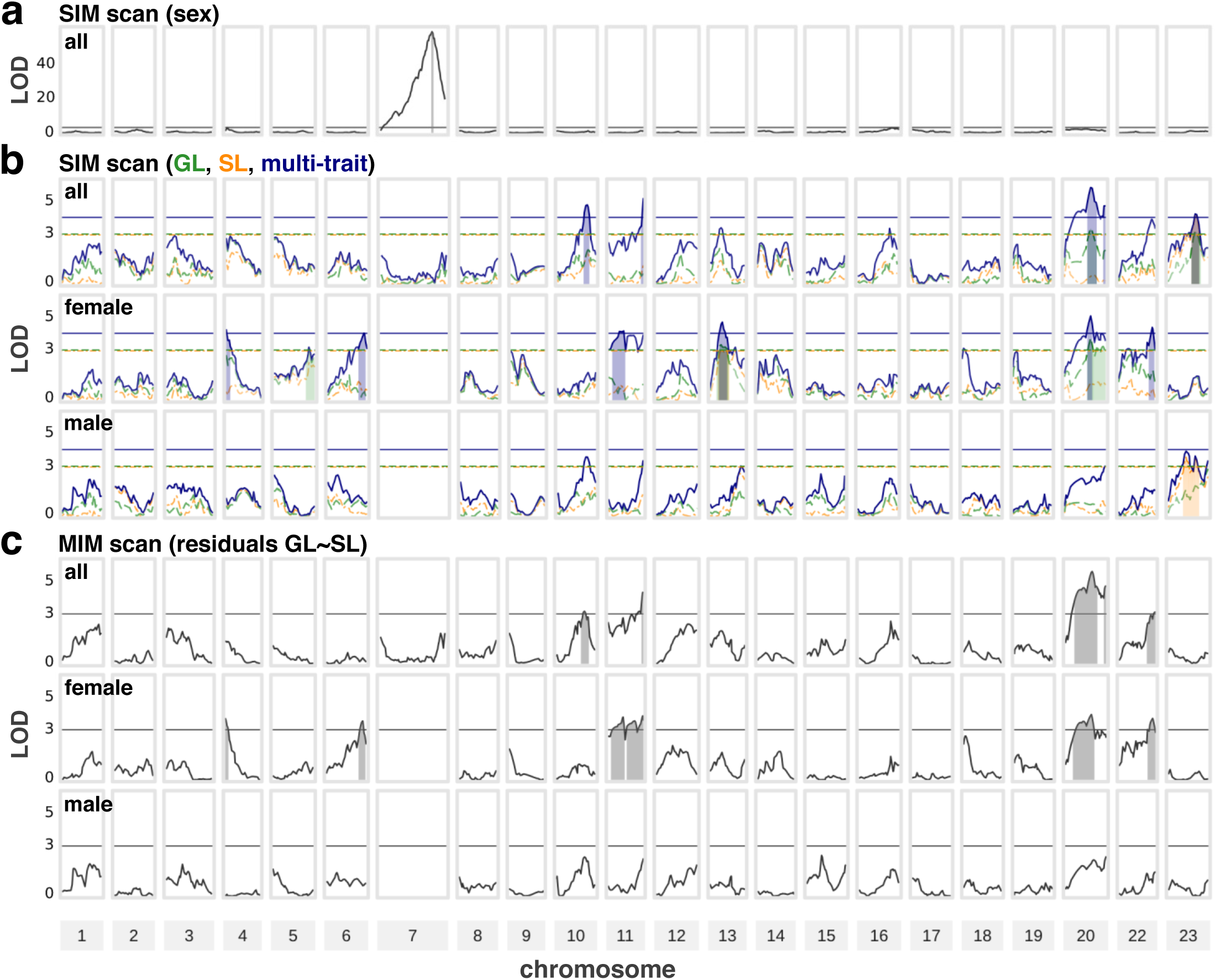
Interval mapping QTL scans for three data subsets. Rows of panels show the LOD score profiles of IM scans performed, plotted across the 22 linkage groups in the genetic map. a) IM scan of sex identifies known XY locus on chromosome 7. b) Multi-trait IM scans performed across three different data subsets (orange lines and shading: SL scans; green: GL scans; dark blue: joint SL-GL scans). c) IM scans of the GL∼SL model residuals (black lines and shading) performed across three data subsets. In b) and c), Horizontal lines indicate significance thresholds color-coded by the trait they represent. Shaded areas represent genetic intervals defined by a drop of 1 LOD from any peak above the significance threshold, and are color- coded by trait. The sex chromosome 7 is omitted from the sex-subsetted scans.

We identified the omnivore *Metriaclima mbenjii* and carnivore *Aulonocara koningsi* to serve as parental species for the F2 cross based on key criteria, including the ability to produce viable and fertile offspring via *in vitro* fertilization, dietary adaptation spanning two trophic levels, and more pronounced differences in gut length (1.89 fold) versus adult body size (1.16 fold) (Fig. 1, a, c, d, and e). The lab lines of the two species were both derived from the same location in Lake Malawi, Mbenji Island, which minimizes potential evolved differences associated with the gut due to environmental factors other than diet. Phenotypically, the F2 hybrid population (*N* = 495) is intermediate to the carnivore and omnivore species in the comparative intestine length survey, with variance spanning the ranges of their parental species (Fig. 1 a). Phenotypic sex was determined for F2 individuals (see Methods) and only those 448 F2 individuals with clear sex information (218 female, 230 male; 1:1.03 female-to-male ratio) were used in subsequent analyses.

Exploratory data analysis of standard body length (SL) and gut length (GL) was performed on the data from the F2 hybrid cross, and data from individuals of the parental species. The mean SL and GL were significatively higher in *M. mbenjii* (SL: *M* = 40.18 mm, *SE* = 0.75 mm; GL: *M* = 78.72 mm, *SE* = 3.31 mm) than in *A. koningsi* (SL: *M* = 34.6 mm, *SE* = 1.24 mm; GL: *M* = 41.62 mm, *SE* = 1.92 mm) with GL showing a greater difference between the parental species compared to SL (Fig. 1, c and d; Welch two sample t-test SL: *t*(14.842) = - 3.8502, *p* = 0.0016; GL: *t*(14.438) = -9.6878, *p* = 1.058e-07). The mean SL and GL of the F2 hybrids (SL: *M* = 37.41 mm, *SE* = 0.27 mm; GL: *M* = 64.69 mm, *SE* = 0.7 mm) had intermediate values to the mean SL and GL of the parental species *M. mbenjii* and *A. koningsi* (Fig. 1, c and d). The linear regression of the model *GL∼SL* showed a positive association between SL and GL (Fig. 1e; *β* = 2.13479, *p* < 2e-16, *R^2^* = 0.6751). There were significant differences by sex in SL and GL in the F2 population (Fig. 1 c and d; Welch two sample t-test SL: *t*(444.89) = - 4.3202, *p* = 1.924e-05; GL: *t*(441.62) = -2.9949, *p* = 0.0029) with males having higher means for both traits relative to females. However, the residuals of the linear regression *GL∼SL* in the F2 population did not show differences by sex (Fig. 1f; Welch two sample t-test *t*(433.92) = 0.91375; *p* = 0.3614). Differences by sex were not tested in the parental species samples due to small sample sizes. The *GL∼SL* residual values of the F2 population span nearly all of the variance present in the multi-species panel, with only eight herbivore individuals having values outside of the range of the F2 hybrids (Fig. 1a). The F2 population also forms a statistically distinct group to the multi-species panel collapsed by trophic level, with mean residuals intermediate to carnivores versus herbivores and omnivores (Fig. 1b; one-way ANOVA, F = 60.8, p < 0.0001; Tukey’s HSD test).

### Mapping strategy

Genetic marker information for the F2 individuals was produced using a ddRADseq strategy. These markers were used to build a genetic map composed of 474 markers distributed across 22 linkage groups, with a length of 1162.4 cM and an average spacing of 2.6 cM. We used this linkage map for subsequent analyses (Supplementary Fig. 1). Below, we describe a multifaceted mapping strategy to identify the number, direction, and magnitude of the QTL contributing to the genetic variance in gut length, including examination of potential epistasis and QTL-by-sex interactions.

### Interval Mapping Analysis

We performed Interval Mapping (IM) as an initial strategy to infer the number of QTL underlying gut length variability, as well as the direction and magnitude of their additive and dominance effects. Because GL is correlated to SL, we used two approaches to account for allometry: 1) a multi-trait mapping approach (Jiang and Zeng, 1995) that consists of mapping SL, GL, and the joint trait SL-GL, and 2) the mapping of the residuals of the model *GL∼SL*. While the F2 population does not show significant differences in GL by sex after controlling for SL (Fig. 1e), previous studies within single species of East African cichlids suggest sexual dimorphism in gut biology (Faber-Hammond et al. 2019, Moore et al. 2022). We thus tested for QTL-by-sex interactions or sex-specific QTL in the genetic architecture of GL by performing the two mapping approaches above on three subsets of the data: 1) all individuals, 2) only female individuals, and 3) only male individuals. Hence, we conducted a total of 12 IM scans resulting from the combination of four traits (SL, GL, joint SL-GL trait, and residuals) and three data subsets (all individuals, males, and females) (Fig. 2b and c). We also conducted a single IM scan for sex (Fig. 2a), confirming the presence of a previously characterized XY locus on chromosome 7 (Ser, 2010) which acts as the sole sex determination locus in this cross.

Our IM analysis identified candidate QTL for at least one of the four traits on nine chromosomes (4, 5, 6, 10, 11, 13, 20, 22, and 23). We found multiple QTL peaks on three of those chromosomes (11, 20, and 23; Fig. 2, Supplementary Table 3); noting the caveat that it can be challenging to discern whether there are multiple QTL on a single chromosome with IM. From these nine chromosomes, we selected a single peak per chromosome and examined their phenotypic effects for SL, GL and residuals. The majority of these candidate QTL exhibited negative additive effects, meaning that the alleles from the *M. mbenjii* (omnivore, long gut) parental species are associated with longer gut lengths and body sizes, and the alleles of the *A. koningsi* (carnivore, short gut) parental species are associated with shorter guts and body sizes, matching expectations. Furthermore, the candidate QTL that were detected in both the GL and SL individual scans had the same direction of additive and dominance effects, as is expected in correlated traits. For example, the candidate QTL detected on chromosomes 5 and 23 are likely body size QTL that also impact GL via body scaling rather than gut-specific action, given the lack of significance at these loci in the scans of residuals (Fig. 3, Supplementary Table 3).

**Fig. 3.**
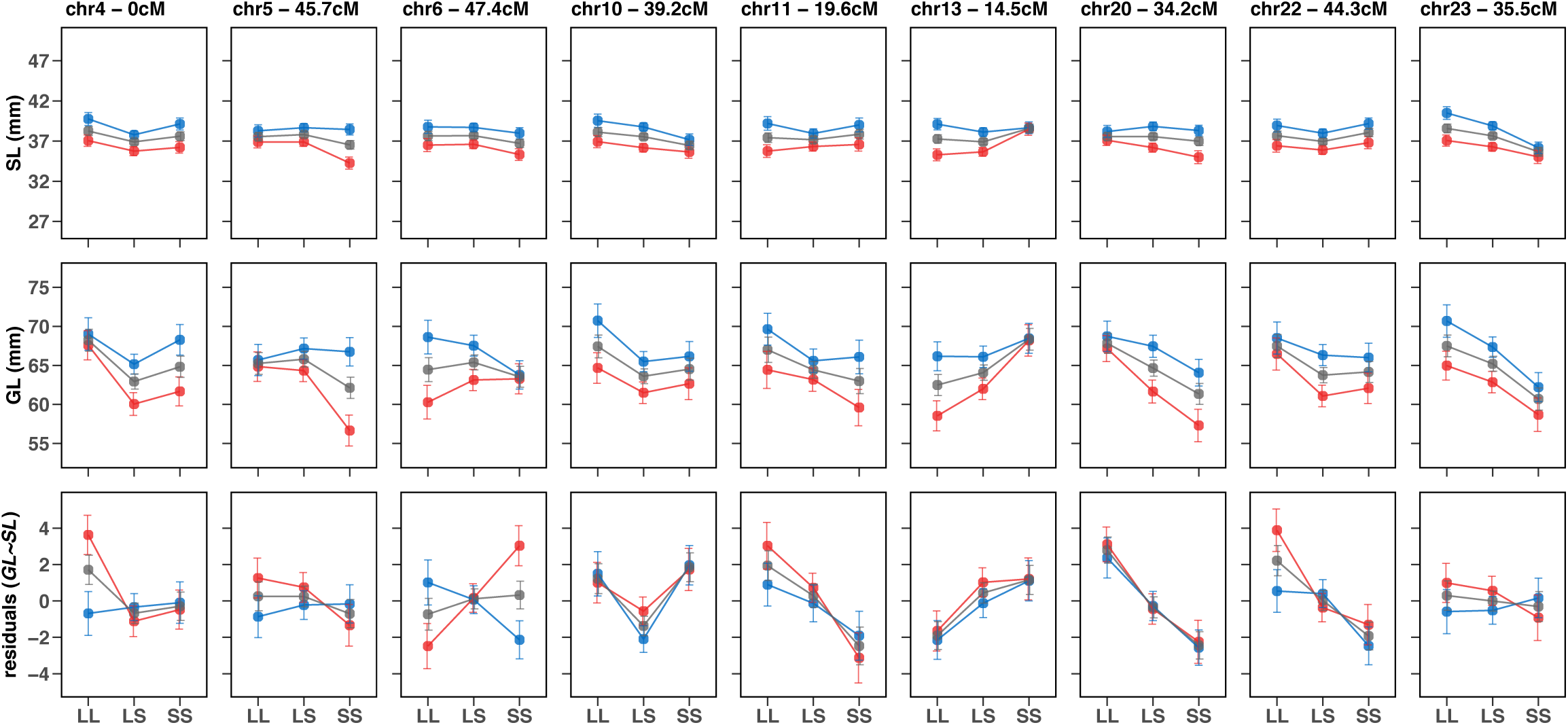
Phenotypic effect plots of nine candidate QTL identified through IM, organized by QTL position (columns) and traits (rows). Each panel shows phenotype values in the F2 hybrids at each candidate QTL by genotype of its most proximate genetic marker (map location at top). Genotypes are labeled based on the parental species they descend from (L allele: *M. mbenji*, long gut species; S allele: *A. koningsi*, short gut species). Effects for the three data subsets are plotted separately: all individuals (gray), females (red), and males (blue). All QTL-trait-data subset combinations are shown, including ones that did not reach the significance threshold.

**Fig 4.**
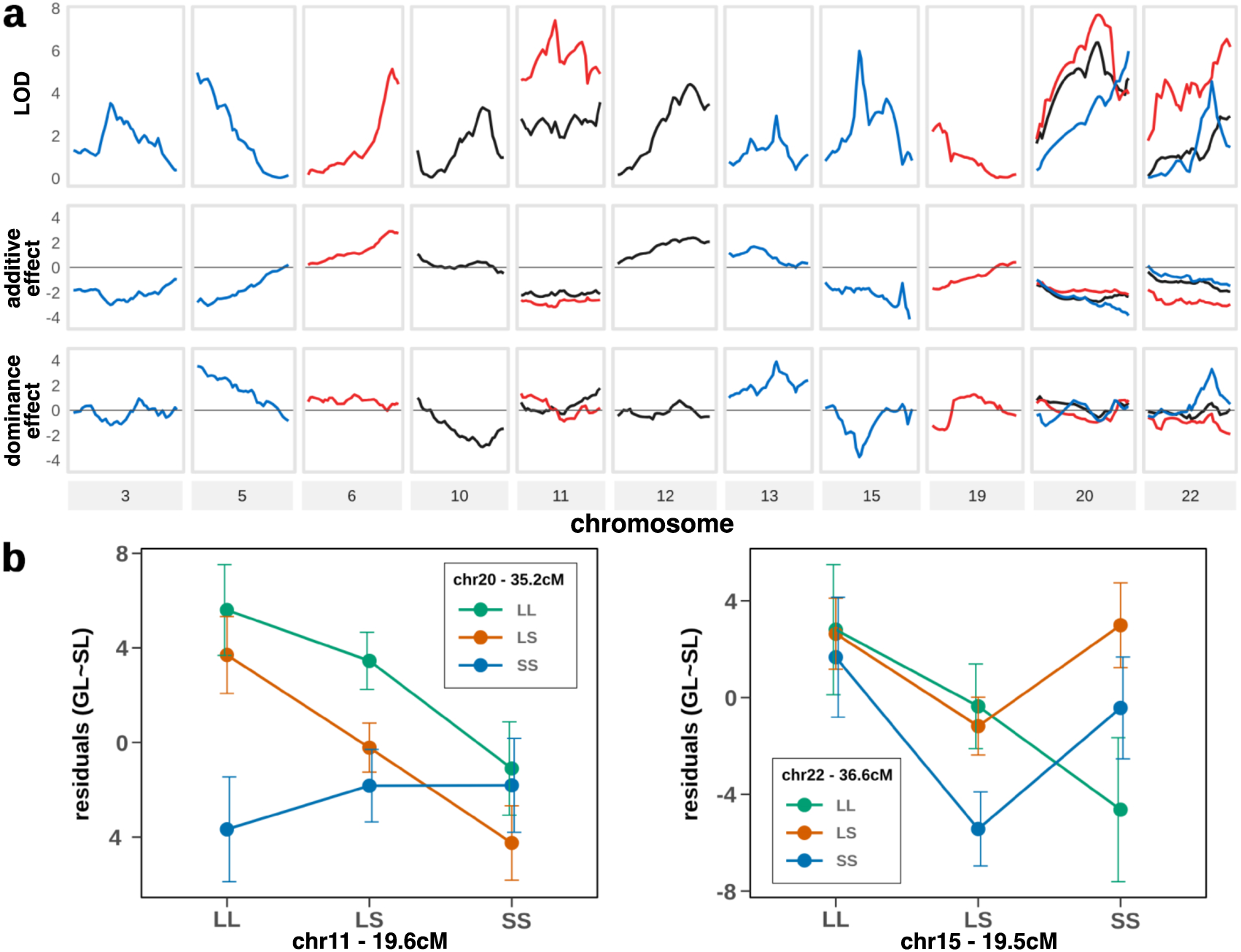
Multiple interval mapping scans of residuals including additive and dominance effect plots. a) Rows of panels show from top to bottom: the LOD score profile, additive effects, and dominance effects of the QTL peaks identified in the residuals MIM analyses, plotted across the chromosomes with significant LOD scores. Line colors represent the data subset used in the analysis where each peak was detected (all individuals, black; females, red; males, blue). The horizontal lines in the additive and dominance effects plots indicate where effects equal zero (no effect) to facilitate interpretation. A negative effect value indicates the expected direction of phenotypic values based on parental genotypes and trait values. b) Phenotypic effect plots of two epistatic interactions identified through MIM. Left, a female-specific epistatic interaction between chromosome 11 and chromosome 20; and right, a male-specific epistatic interaction between chromosome 15 and chromosome 22. Genotypes are labeled based on the parental species they descend from (L allele: *M. mbenjii,* long gut species; S allele: *A. koningsi*, short gut species).

Most QTL were detected in the joint SL-GL and residuals trait scans. Peaks found in these scans can be driven by the presence of QTL with discordant effects between SL and GL, or QTL that only impact one of the two traits. In the case of the SL-GL joint scan, a marked increase in the LOD score of the joint scan is expected relative to the individual scans of SL and GL when these two traits are decorrelated (Jiang and Zeng, 1995). Similarly, the residuals scan should exclude any peaks driven by allometry because the phenotypic variance of gut length that is explained by body size has been removed through linear regression. The difference between the SL-GL joint scan and the residuals scan is that the former is also able to identify QTL that may be specific to SL. Indeed, there are three QTL (on chromosomes 5, 13 and 23) that were detected by the GL, SL, or joint SL-GL scans and were absent in the residuals scans. The female QTL on chromosome 13 is the only one of these three QTL that displays a significant peak that is higher in the joint SL-GL scan compared to the individual SL and GL scans, suggesting that there may be uncoupling of SL and GL at this locus. Given the patterns observed in the effect plots at this locus (Figs. 2 and 3), it is possible that there are SL-specific and GL-specific QTL in close proximity to each other in this region, but only the SL-specific QTL reaches significance. Notably, the remaining six QTL, found on chromosomes 4, 6, 10, 11, 20, and 22, all seem to display uncoupling between SL and GL and are likely GL-specific QTL. For instance, the GL and joint SL-GL scans in the all individuals and females subsets display significant peaks on chromosome 20, while the SL scans do not display peaks on this chromosome. Additionally, in chromosome 20, the LOD scores are noticeably higher in the joint SL-GL peaks compared to the GL peaks, and the residuals scans recapitulate the results observed in the joint SL-GL scans. Similarly, chromosome 10 displays joint SL-GL and residuals peaks in the all individuals subset. None of the individual trait scans (SL or GL) display a significant peak at this locus, suggesting that this QTL is more easily detected when the interplay of SL and GL is considered using the joint SL-GL and residuals scans. This observation is further supported by the effect plots of this QTL which show a clearer dominance effect in the residuals than in the SL and GL traits. (Figs. 2 and 3).

Additionally, one of the prominent patterns in the IM analyses is the pervasive sex differences observed in the QTL scans and phenotypic effects. One striking example of differences between sexes is found on chromosome 6, where a QTL was detected solely in the females joint SL-GL and residuals scans, suggesting that this is a GL-specific QTL (i.e., not impacted by allometry) that also interacts with sex or is specific to females. This interpretation is further supported by the phenotypic effect plots which show a negative additive effect in the residuals trait for males at this position, while females display a positive additive effect. Other loci where the additive effects of females and males seem to differ include chromosome 4 (GL and residuals), chromosome 5 (GL), chromosome 13 (SL and GL), and chromosome 22 (residuals) (Figs. 2 and 3). Sex-by-genotype effects at many of these QTL suggest the presence of alleles modulating the degree of sexual dimorphism in the traits examined. In the case of the chromosome 13 QTL, individuals homozygous for the long gut species allele display sex dimorphism in SL and GL, while those homozygous for the short gut species allele do not; however, there is no sex dimorphism in the residuals trait where allometry is taken into account (Fig. 3). Alternatively, at the chromosome 4 QTL, sex dimorphism is present for SL in all genotypes, but appears to be lacking for GL only in those individuals homozygous for the long gut species allele; the net outcome is that only these homozygotes show sexual dimorphism in the residuals trait (Fig. 3). These scenarios suggest evolved genetic differences tuning sexual dimorphism within species for individual traits. Here, we hypothesize that the chromosome 13 QTL may modulate sexual dimorphism in overall body size, while the chromosome 4 QTL may modulate sexual dimorphism in gut length.

There are a few QTL that are more challenging to interpret, such as the QTL on chromosomes 11 and 20. These QTL display similar additive and dominance effects in both sexes in the residuals trait (Fig. 3) but the male residuals scan does not reach significance at these positions (Fig. 2). When looking at the chromosome-level view of the additive and dominance effects of chromosomes 11 and 20 (Supplementary Fig. 3), we can observe that the dominance effects differ substantially between females and males, suggesting that there might be QTL-by-sex interactions or multiple sex-specific QTL within these chromosomes at different positions. Apart from the differences in phenotypic effects, it is worth noting that only a single QTL -the standard length QTL on chromosome 23- was detected in the male subset scans; in contrast, 7 QTL were detected in the female subset scans (Figs. 2 and 3). Though this difference may be in part due to the limited power of the female and male IM scans, this result may also indicate that the genetic architecture of gut length of males and females have minimal overlap. Moreover, IM scans do not control for variance explained by background QTL or epistatic interactions, which may differently impact the female and male scans given the presence of QTL-by-sex interactions or sex-specific QTL.

Overall, concordance among the various IM scans suggests that several of the candidate QTL are specific to gut length and are not a consequence of unintended mapping of allometry. In addition, our results suggest that multiple candidate QTL identified through IM involve QTL-by-sex interactions or sex-specific QTL.

### Multiple Interval Mapping

In order to further characterize the genetic architecture of SL, GL, and residuals, we performed Multiple Interval Mapping (MIM) (Kao et al., 1999), a model selection method that enables us to estimate the additive and dominance effects of QTL while accounting for the residual variance of background QTL, as well as to test for epistatic interactions between QTL. As before, we performed our analyses in three different subsets of the data; conducting a total of 9 MIM analyses resulting from the combination of three traits (SL, GL, and residuals) and three data subsets (all individuals, males, and females; Table 1 and Supplementary Table 4). We did not perform the joint analysis of SL-GL as in the IM analysis because multi-trait MIM analysis has not been implemented in standard QTL mapping tools (Broman et al., 2003; Wang et al., 2012), and because IM of residuals of the model *GL∼SL* produced results largely concordant with the joint analysis of SL-GL.

**Table 1.**
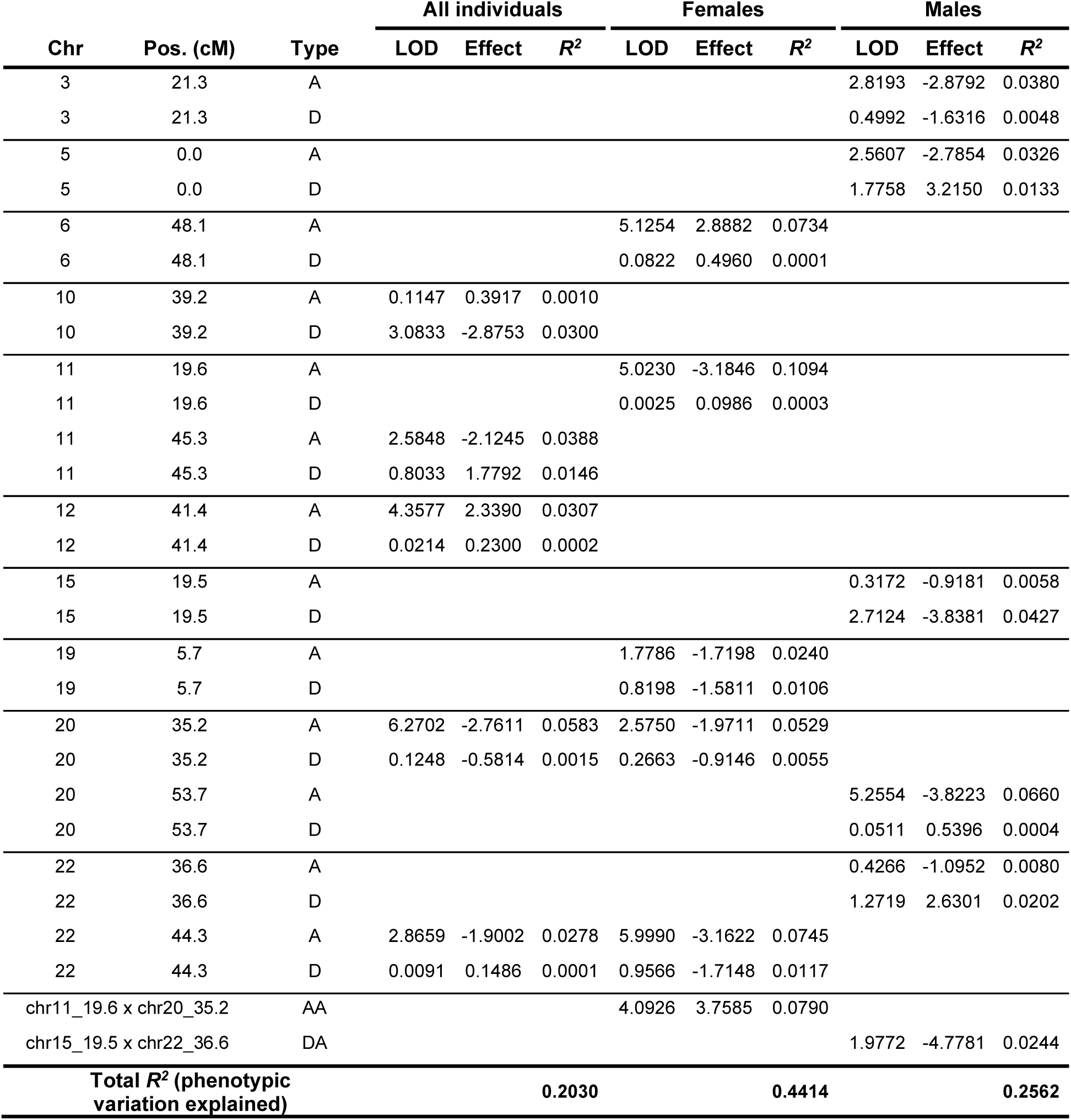
QTL positions and effects from multiple interval mapping of residuals of GL∼SL.

The results of our MIM analyses align with the results obtained through IM, as eight out of the nine candidate QTL identified in the IM analyses were present in at least one of the final MIM models. Several new main effect QTL were detected through our MIM analyses across all traits, highlighting the increased capacity to detect QTL when accounting for background QTL variance. For instance, the residuals analysis in the all individuals dataset detected a new QTL on chromosome 12, the residuals analysis in females detected a new QTL on chromosome 19, and the residuals analysis in males detected new QTL on chromosomes 3, 5, 15, 20, and 22 (Table 1, Fig. 4a). Similarly, the GL analysis in females detected a new QTL on chromosome 14, and the SL analysis in males detected a new QTL in chromosomes 2 and 13 (Supplementary Fig. 4, Supplementary Table 4). In addition to main effect QTLs, three epistatic interactions were identified in the residuals and GL MIM analyses (Table 1, Fig. 4b, and Supplementary Table 4). It is worth noting that the final models of the residuals MIM analyses have overall a higher number of detected QTL as well as percentage of variance explained (approximately 20 to 44 percent, depending on the data subset; Table 1), likely because this is the only trait that accounts for allometry, and because there are more distinct GL differences between the parental species compared to SL differences (Fig. 1). Because of this, we subsequently focused on the interpretation of the MIM analyses of the residuals phenotype (Table 1).

Across all datasets, the residuals MIM analyses showed a majority of additive effects with a negative direction (Fig. 4a), matching expectations based on the parental genotypes. Three QTL (on chromosomes 10, 11, and 12) were detected exclusively in the all individuals analysis. These QTL may be present in both males and females but were unable to be detected in the female or male analyses due to the reduced power of these subsets. Only chromosomes 20 and 22 had QTL across all three data subsets with similar direction of additive effects, though the peaks of the female and male QTL were detected more than 10 cM apart from each other. It is possible that the signals detected in females and males in these two chromosomes are a consequence of shared QTL between females and males; however, given the differences between sexes in additive and dominance effects observed throughout chromosomes 20 and 22 (Fig. 4a), it is also possible that these chromosomes contain multiple linked QTL, some of which may or may not be sex-specific. In contrast, none of the remaining QTL (chromosomes 3, 5, 6, 11, and 15) were identified in both the female and male analyses, supporting our previous observations of marked differences between the genetic architecture of males and females.

Additionally, two sex-specific QTL-by-QTL epistatic interactions were detected in the residuals scans: 1) an additive-by-additive interaction between chromosomes 11 and 20, detected in the residuals analysis of the female data subset; and 2) an additive-by-dominant interaction between chromosomes 15 and 22, detected in the residuals analysis of the male data subset (Table 1, Fig. 4 b). Notably, the chromosome 11 and 20 epistatic interaction involves the two QTL that produced similar phenotypic effects but different mapping results by sex in the IM analysis (see above).

Given that our MIM analyses were performed using a relatively strict statistical threshold that could inadvertently suggest significance in one sex but not the other due to small differences in statistical power, we replicated the female and male SL and residuals MIM analyses using a more lenient statistical threshold to corroborate that our observations are not an artifact of the chosen statistical threshold (see Methods; Supplementary Table 5). The results of the lenient-threshold analyses detected a larger number of main effect QTL and QTL-by-QTL interaction terms, as expected. Some of the newly detected terms also supported some of our predictions; for instance, the QTL previously detected in the all individuals subset in chromosome 10 reached significance in the male subset, and a new female residuals QTL was detected in chromosome 13 in close proximity to the previously detected SL-specific QTL. However, the lenient-threshold-based analyses also showed marked differences between the sets of QTL detected in females and males, providing continued support for the presence of pervasive sex-specific QTL in the residuals genetic architecture (Supplementary Table 5).

Overall, our MIM results showed a large overlap with our IM analysis, and MIM allowed us to refine further the location, number, and magnitude of the effects of the QTL underlying SL, GL, and the residuals of the model *GL∼SL* in our F2 mapping cross. The differences in the number of QTL detected across traits and the percentage of variance explained by our final models suggest that our cross is better suited for mapping the genetic architecture of gut length than the genetic architecture of body size, a result that matches our observations in the phenotypic analysis of the parental species. The final models of the residuals MIM analyses also confirm the presence of abundant sex-specific components in the genetic architecture of the residuals, including sex-limited epistatic interactions, indicating that the genetic architecture of the residuals is sex-specific.

## Discussion

By comparing multiple species from the Lake Malawi cichlid radiation, we have shown that this group of species recapitulates a broad gastrointestinal trend in vertebrates: herbivore and omnivore species tend to have longer guts compared to carnivore species. Moreover, by raising these species under controlled dietary and laboratory conditions, and performing genetic mapping in a hybrid cross between two of these species, we have demonstrated that observed differences in gut length have a significant, evolved genetic basis. The results of our multifaceted mapping strategy suggest that the genetic architecture of evolved gut length differences in Malawi cichlids is polygenic and complex, involving epistatic interactions, pervasive sex-by-genotype effects, and variants impacting the degree of sexual dimorphism in traits. Indeed, few QTL were shared between sexes even under lenient statistical conditions, suggesting largely distinct, sex-specific genetic architectures for gut length.

### Sexual conflict and the gut

Scenarios where males and females have different fitness optima for a particular trait can lead to sexual conflicts, which in turn can be resolved through evolution of sexual dimorphism. Such resolution involves evolution of genetic variants that constrain the phenotypic effects of genotypes to a single sex, in other words, through the evolution of sex-specific genetic architectures (Mank, 2017). Our results show limited overlap in the MIM QTL detected in female and male data subsets in the three traits analyzed (residuals, GL, or SL) (Table 1, Supplementary Tables 3, 4, and 5), and some of the QTL involved appear to modulate the degree of sexual dimorphism in SL, GL, or the allometric relationship between the two (Fig. 3). The marked differences in the genetic architecture of males and females strongly suggest that the genetic architecture of gut length (and possibly standard length) is sex-specific, and thus we conclude that gut length in Malawi cichlids is a trait with a history of sexual conflicts that have been resolved through the evolution of sex-specific genetic architectures. Notably, in a recent mapping study in a hybrid cross between cave and surface populations of the fish *Astyanax mexicanus*, two of the three hindgut length QTL identified had sex-specific effects (Riddle et al., 2021), suggesting that sex-specific genetic architectures for gastrointestinal traits may be a broad phenomenon. A mapping experiment in a hybrid cross of Lake Victoria cichlids identified a single intestine length QTL, but the study did not include sex-specific mapping (Feller et al., 2022); the QTL mapped in the Victorian cross did not overlap with any of the QTL we identified here. In our study, only 5 out of 13 main effect QTL in the more conservative MIM analysis would have been identified in the residuals MIM analysis if sex had not been considered. We thus suggest careful consideration of sex-associated biology in forward genetics studies, even where the phenotype itself is not overtly sexually dimorphic, as is the case here when comparing the *GL∼SL* residuals in our F2 population by sex. That is, sexually monomorphic traits may require sexually dimorphic genetic architectures to respond to sex-specific differences in development and physiology, and sex-blind mapping strategies may miss important genotype-phenotype associations.

Sexual dimorphism is a hallmark of Lake Malawi cichlids with obvious examples including nuptial pigmentation and territorial behavior related to mating (Konings, 2007; Roberts et al., 2009). Recent studies comparing traits within single cichlid species have identified sex-associated differences in gut microbiota and intestinal length, suggesting the presence of sex-specific fitness optima for gastrointestinal traits as well (Faber-Hammond et al., 2019; Moore et al., 2022). Sexually dimorphic life history differences related to territoriality and parental investment provide compelling potential explanations for sexual conflicts associated with gut length. Most notably, Lake Malawi cichlids are maternal mouthbrooders, a parenting strategy which requires females to undergo cyclical and prolonged periods of starvation (roughly three weeks per brood) while incubating developing offspring in their mouth until they become free-swimming fry. Additionally, males of many species including *M. mbenjii* are highly territorial and tend to remain in a very limited space, which may pose restrictions in the diversity and quantity of food sources available for foraging (Ribbink, 1983). Thus, limitations in food access for males may be more constant and less severe compared to females, who undergo prolonged, alternating periods of fasting and unrestricted foraging. The idea that differences in parental care lead to sex dimorphism in trophic traits is not novel. In Tanganyikan cichlids, another trophic trait, gill raker length, is sexually dimorphic in uniparental mouthbrooders but not biparental mouthbrooders, providing strong support that sexual dimorphism in reproductive strategies can lead to sexual dimorphism in trophic traits (Ronco et al., 2019). Body size is also sexually dimorphic to varying degrees among Lake Malawi cichlids, with males generally being larger in size (Genner and Turner, 2005). Given the interactive relationship between body size and gut length, sex-specific genetic architectures may have co-evolved to counter maladaptive outcomes in gut length for one or both sexes arising from evolution of sexual dimorphism in body size. Finally, general sex differences in development and physiology may require sex-specific genetic architectures to produce similar trait outcomes, that is, different genetic architectures may be necessary to mitigate the effects of different sex "environments" to produce non-sexually dimorphic trait values. All of these scenarios are ultimately linked to the evolutionary trade-off between maintaining a longer gut that allows for increased absorption of nutrients, versus the high energy requirements of maintaining the gut tissue (Cant, 1996). We note that cichlid species radiations provide a powerful comparative model to explore the above hypotheses, for example by comparing species with different parental strategies (e.g., comparisons among paternal, maternal, and biparental mouthbrooders), or comparing species with varying levels of territoriality or sexual dimorphism in body size.

### Implications of epistatic interactions in the genetic architecture of gut length

Previous studies have shown that gene-by-gene interactions, though sometimes overlooked, are a prevalent component of the genetic architecture of quantitative traits (Mackay, 2014). In this study, we detected the presence of two epistatic interactions in the residuals MIM analyses. The first one is an additive-by-additive interaction found in the genetic architecture of the residuals in females, involving QTL on chromosomes 11 and 20 (Fig. 4b). In this interaction, the negative additive effect of the QTL on either chromosome 11 or 20 is masked in the presence of an SS genotype at the other locus. This reciprocal suppressing epistatic interaction may provide a canalization mechanism in the wild, where an *A. koningsi* SS genotype at either locus contributes to a shorter-gut phenotype despite variation at the other locus. Genetic canalization produces invariability in a phenotype despite the presence of genetic variants capable of modifying such phenotype (Waddington, 1942), that is, it can help maintain cryptic genetic variation. Canalization may support rapid evolution by providing a reservoir of potentially adaptive variation that can be quickly released from its cryptic state for selection to act upon, or by buffering maladaptive variation that would otherwise reduce fitness (Gibson and Dworkin, 2004). Introgression and interspecific hybridization have likely supported the rapid radiation of Malawi cichlid species (Malinsky, 2018), perhaps in part through the masking and unmasking of genetic variation through epistasis, with alleles being naturally selected in varied genetic backgrounds through hybridization events. Experimental crosses in cichlids support this scenario, for example where hybrids have more extreme variation in body morphology than found among parental populations, possibly due to unmasked cryptic variation (Nichols et al., 2015; DeLorenzo et al., 2023). Our results suggest specific predictions; for example, that chromosome 20 variation would be subject to selection on a chromosome 11 LL-genotype background, potentially leading to fixation of one chromosome 20 allele; on the other hand, chromosome 20 variation would be cryptic on a chromosome 11 SS-genotype background and would be subject to genetic drift, potentially maintaining chromosome 20 variation.

The second epistatic interaction is a dominant-by-additive interaction found in the genetic architecture of the residuals in males, involving QTL on chromosomes 15 and 22 (Fig. 4b). In this interaction, the QTL on chromosome 15 has an expected negative additive effect only in the presence of the LL genotype at chromosome 22, while an LL genotype at chromosome 15 masks the effect of the QTL on chromosome 22. Furthermore, the chromosome 15 QTL displays underdominance in the presence of an SS or SL genotype on chromosome 22. While this interaction is complex and less immediately interpretable than the one described above, it reveals the unpredictable phenotypic outcomes that epistasis can produce for complex traits, particularly in hybrids.

## Conclusion

Given the limited understanding of the genetic basis of the morphological and physiological gastrointestinal diversity observed across animals, the results here and from similar studies are key to understanding how variation in the gastrointestinal system arises at the genome level, and for identifying novel pathways and targets for medical, environmental, and agricultural applications. Our results suggest that sexual dimorphism in the genetic architecture of gut length in cichlids may have evolved as a way to resolve sexual conflicts stemming from differences in reproductive strategies, or general sex differences in development and physiology. We highlight the importance of studying sexual conflict and sexual dimorphism in traits not overtly related to reproduction or sexual fitness, such as gastrointestinal traits, and generally in traits that appear sexually monomorphic. We also find evidence of epistasis and canalization underlying interspecific gut length differences, providing opportunity for maintenance or release of cryptic variation during hybridization, depending on genetic background. The adaptive radiation of Lake Malawi cichlids is marked by regular hybridization and introgression (Malinsky et al., 2018), and complex sex determination systems that uncouple sex-limited and sex-linked trait effects (Moore et al., 2022). Together with the genetic architecture of gut length we describe here, we speculate that the interplay of sexual conflict, hybridization, and epistasis contributed to rapid and ongoing evolution of diverse trophic strategies in Lake Malawi cichlids.

## Acknowledgements

We thank Maggie Cline, Lindsey Gentry, Adam Miranda, Jessica Potts, Megan Russell, Clare Stull, Jodie White, and Briana Williams for their animal husbandry support of the project.

## Data availability statement

Phenotype data is available in Supplementary Tables 6 and 7. Raw sequence data are available at: https://www.ncbi.nlm.nih.gov/bioproject/PRJNA955776

## Conflict of interest

The authors declare no conflicts of interest.

**Supplementary Table 1.** Species tested with assigned trophic levels and statistical comparisons of residuals of the model GL∼SL.

**Supplementary Table 2.** Segregation distortion on Chr 11 by F1 dam. F2 families from one of the two F1 dams lack the *Aulonocara* homozygous genotype on Chr 11.

**Supplementary Table 3.** Full interval mapping results for all traits in all data subsets.

**Supplementary Table 4.** Full multiple interval mapping results for all traits in all data subsets.

**Supplementary Table 5.** Multiple interval mapping results for GL and SL in males and females comparing strict and lenient statistical thresholds.

**Supplementary Table 6.** Phenotypic data of multi-species panel.

**Supplementary Table 7.** Phenotypic data for hybrid individuals with unambiguous sex, used for mapping.

**Supplementary** Figure 1. Graphic depiction of genetic linkage map from marker analysis in F2 hybrids.

**Supplementary** Figure 2. Interval mapping scans of SL including additive and dominance effect plots.

**Supplementary** Figure 3. Interval mapping scans of residuals of GL∼SL including additive and dominance effect plots.

**Supplementary** Figure 4. Multiple interval mapping scans of SL including additive and dominance effect plots.

---

**Supplementary Fig. 1.**
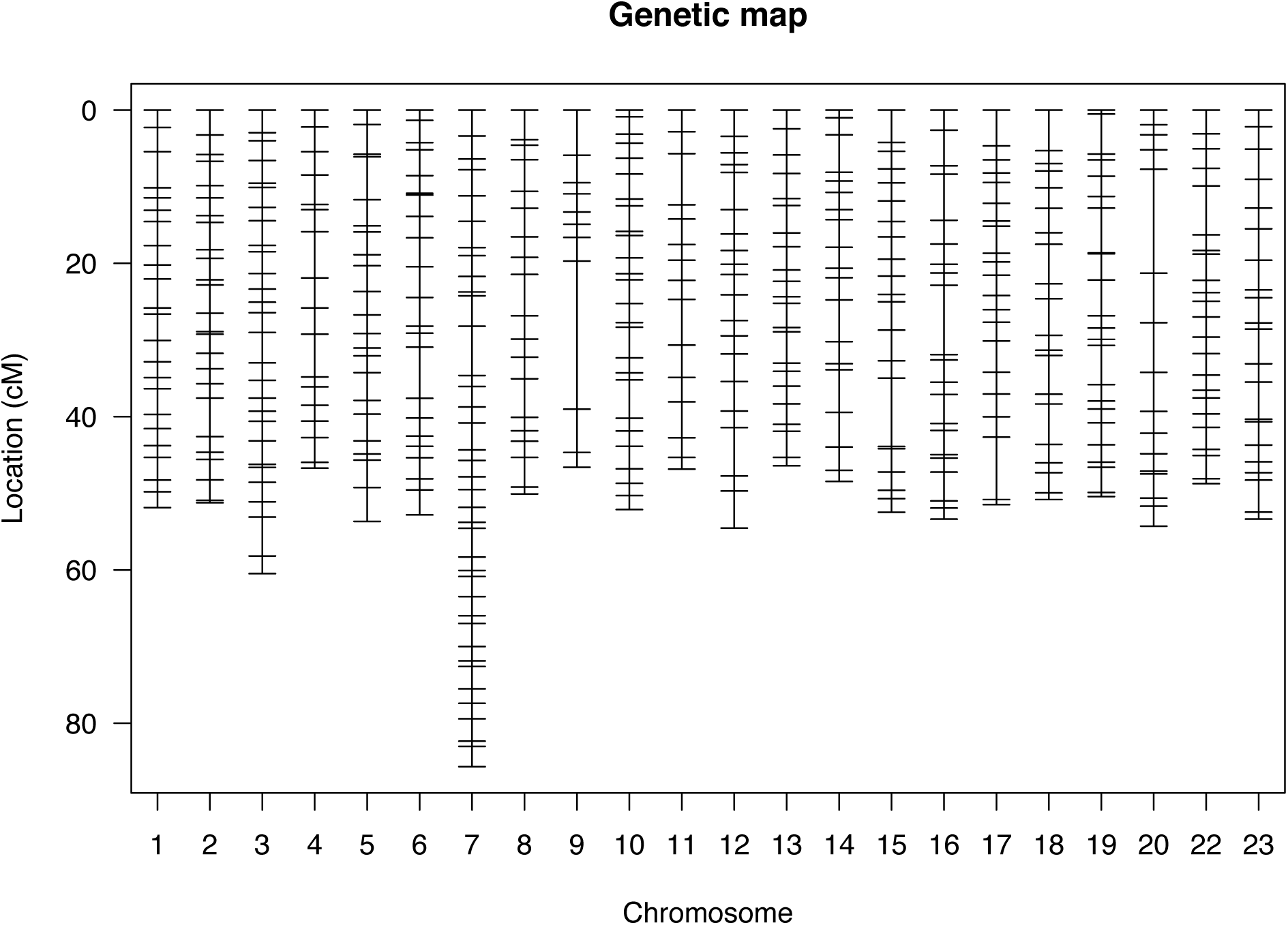
Graphical depiction of genetic linkage map from marker analysis in F2 hybrids. The genetic map consists of 474 markers distributed across 22 linkage groups, matching the number of chromosomes in the cichlid genome. The markers have an average spacing of 2.6 cM, and the map has a total length of 1162.4 cM. Chromosomes are numbered based on established convention for the standard Lake Malawi cichlid karyotype, which skips number 21.

**Supplementary Fig. 2.**
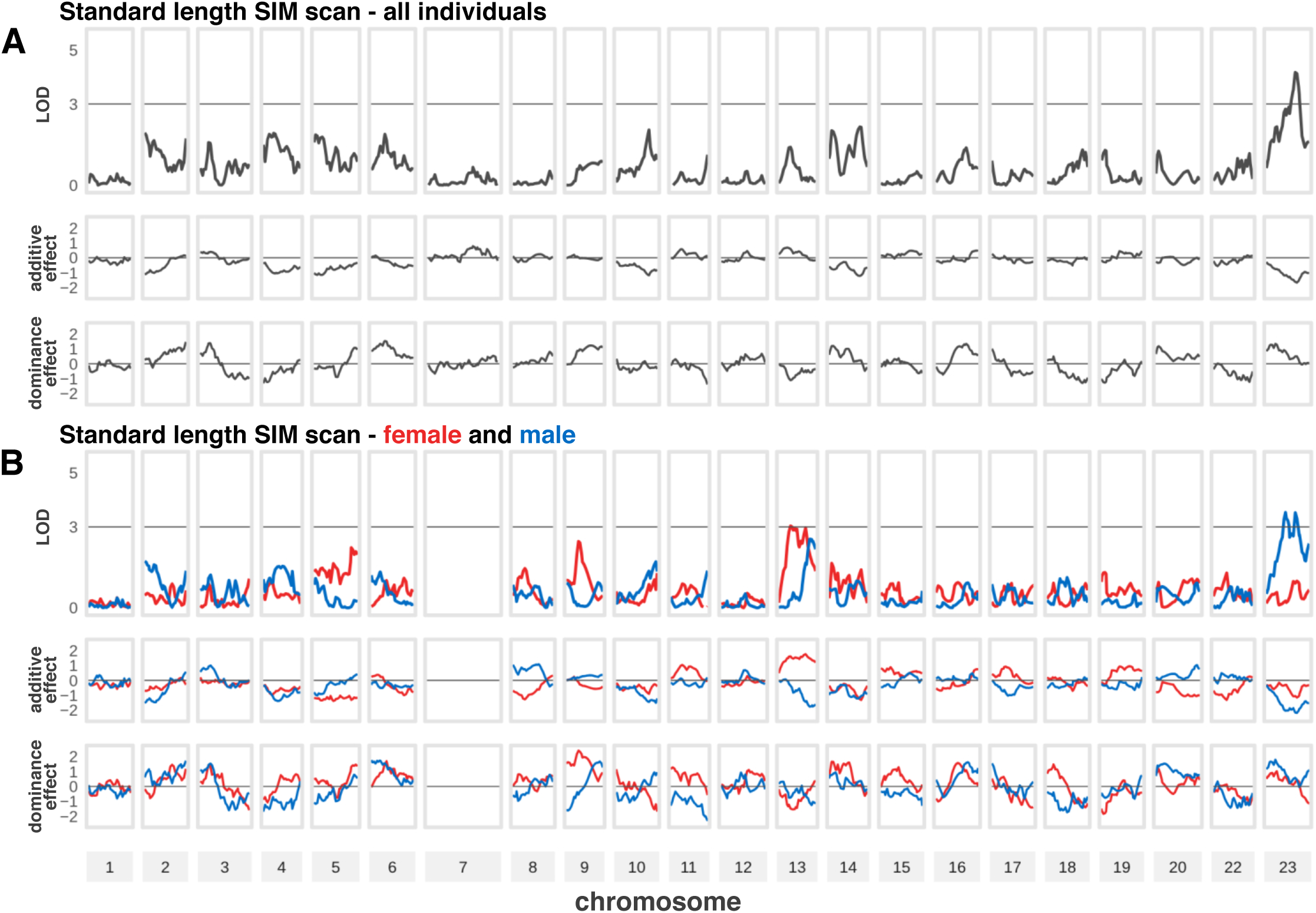
Interval mapping scans of SL including additive and dominance effect plots. Rows of panels show the LOD score profile, additive effects, and dominance effects of IM scans performed, plotted across the 22 linkage groups in the genetic map. a) IM scan of standard length including all individuals (N = 448). b) IM scans of standard length for the female (red lines, N = 218) and male (blue lines, N = 230) data subsets. The horizontal lines in the LOD score profile plots indicate significance thresholds. The horizontal lines in the additive and dominance effects plots indicate where effects equal zero (no effect) to aid interpretation. A negative additive effect value indicates the expected direction of phenotypic values based on parental genotypes and trait values.

**Supplementary Fig. 3.**
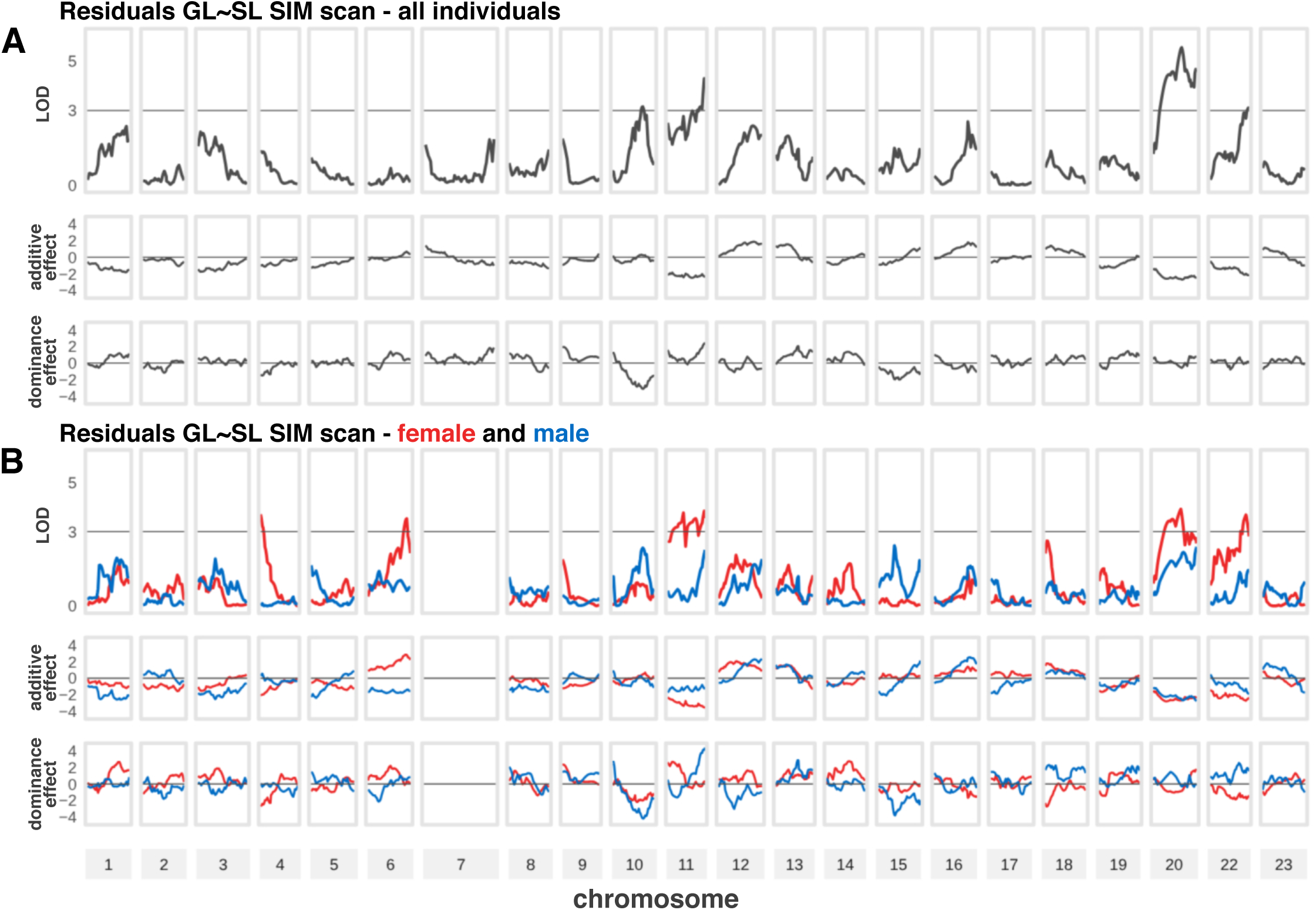
Interval mapping scans of residuals including additive and dominance effect plots. Rows of panels show the LOD score profile, additive effects, and dominance effects of IM scans performed, plotted across the 22 linkage groups in the genetic map. a) IM scan of residuals including all individuals (N = 448). b) IM scans of residuals for the female (red lines, N = 218) and male (blue lines, N = 230) data subsets. The horizontal lines in the LOD score profile plots indicate significance thresholds. The horizontal lines in the additive and dominance effects plots indicate where effects equal zero (no effect) to aid interpretation. A negative additive effect value indicates the expected direction of phenotypic values based on parental genotypes and trait values.

**Supplementary Fig. 4.**
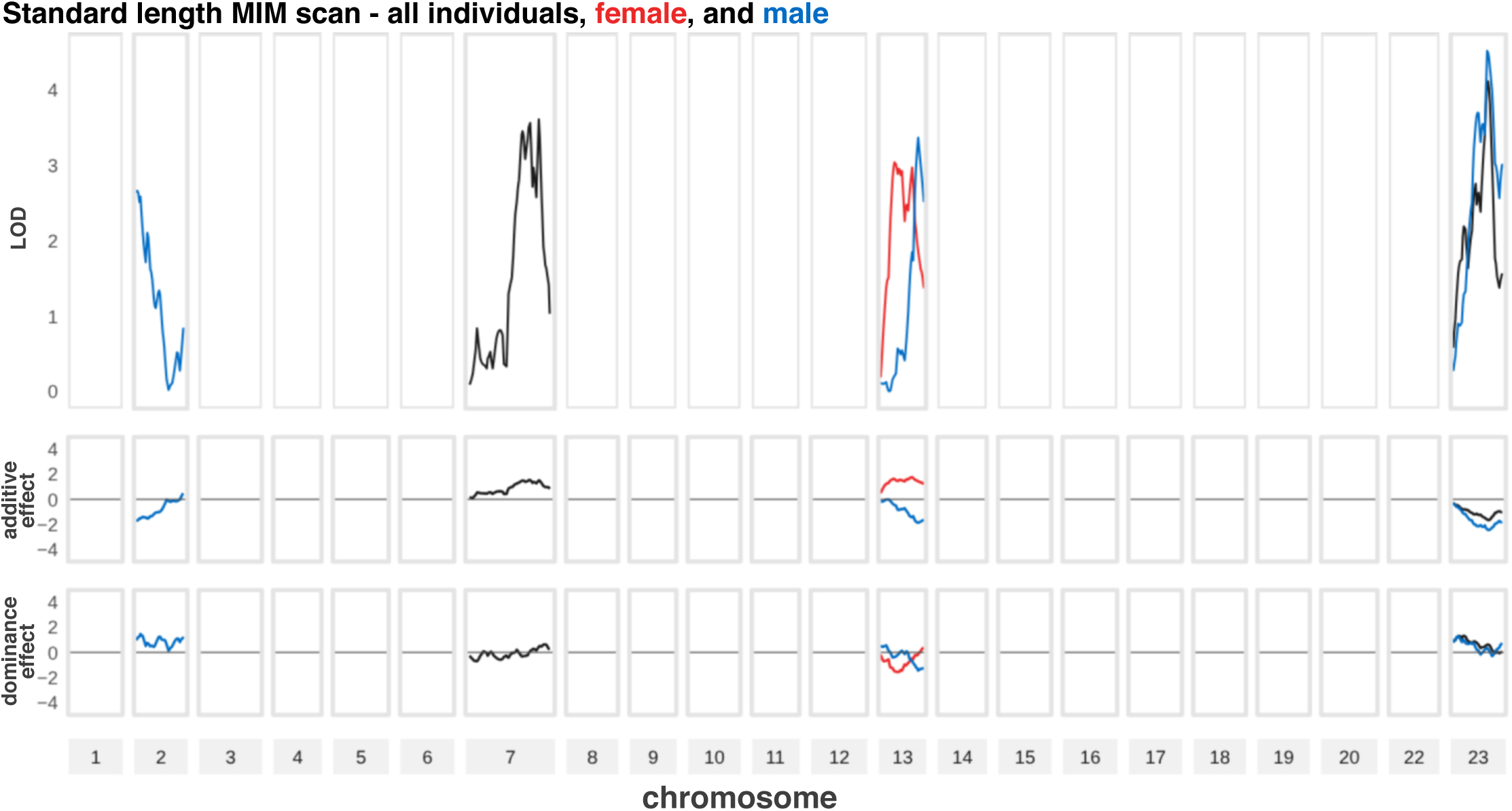
Multiple interval mapping scans of SL including additive and dominance effect plots. Rows of panels show the LOD score profile, additive effects, and dominance effects of the significant QTL peaks identified in the SL MIM analyses, across the 22 linkage groups in the genetic map. Line colors represent the data subset used in the analysis where each peak was detected (all individuals, black; females, red; males, blue). The horizontal lines in the additive and dominance effects plots indicate where effects equal zero (no effect) to aid interpretation. A negative additive effect value indicates the expected direction of phenotypic values based on parental genotypes and trait values. The final model of the all-individuals MIM analysis did not include sex as a covariate; thus, the peak on chromosome 7 likely overlaps the sex determination region and may be associated with sexual dimorphism in body size.

## References

1. Albertson, R. C., Streelman, J. T., & Kocher, T. D. (2003). Directional selection has shaped the oral jaws of Lake Malawi cichlid fishes. Proceedings of the National Academy of Sciences, 100(9), 5252–5257. 10.1073/pnas.0930235100

2. Brawand D, Wagner CE, Li YI, Malinsky M, Keller I, Fan S, Simakov O, Ng AY, Lim ZW, Bezault E, Turner-Maier J, Johnson J, Alcazar R, Noh HJ, Russell P, Aken B, Alföldi J, Amemiya C, Azzouzi N, Baroiller JF, Barloy-Hubler F, Berlin A, Bloomquist R, Carleton KL, Conte MA, D’Cotta H, Eshel O, Gaffney L, Galibert F, Gante HF, Gnerre S, Greuter L, Guyon R, Haddad NS, Haerty W, Harris RM, Hofmann HA, Hourlier T, Hulata G, Jaffe DB, Lara M, Lee AP, MacCallum I, Mwaiko S, Nikaido M, Nishihara H, Ozouf-Costaz C, Penman DJ, Przybylski D, Rakotomanga M, Renn SCP, Ribeiro FJ, Ron M, Salzburger W, Sanchez-Pulido L, Santos ME, Searle S, Sharpe T, Swofford R, Tan FJ, Williams L, Young S, Yin S, Okada N, Kocher TD, Miska EA, Lander ES, Venkatesh B, Fernald RD, Meyer A, Ponting CP, Streelman JT, Lindblad-Toh K, Seehausen O, Di Palma F. The genomic substrate for adaptive radiation in African cichlid fish. Nature. 2014 Sep 18;513(7518):375–381.

3. Broman, K.W., Wu, H., Sen, S. and Churchill, G.A., 2003. R/qtl: QTL mapping in experimental crosses. bioinformatics, 19(7), pp.889–890.

4. Burford Reiskind MO, Coyle K, Daniels HV, Labadie P, Reiskind MH, Roberts NB, Roberts RB, Vargo EL, Schaff J. Development of a universal double- digest RAD sequencing approach for a group of non-model, ecologically and economically important insect and fish taxa. Mol Ecol Resour. 2016; 16(6):1303–1314.

5. Cant, J.P., McBride, B.W. and Croom Jr, W.J., 1996. The regulation of intestinal metabolism and its impact on whole animal energetics. Journal of animal science, 74(10), pp.2541–2553.

6. Catchen, J.M., Amores, A., Hohenlohe, P., Cresko, W. and Postlethwait, J.H., 2011. Stacks: building and genotyping loci de novo from short-read sequences. G3: Genes| genomes| genetics, 1(3), pp.171–182.

7. Conte, M.A., Joshi, R., Moore, E.C., Nandamuri, S.P., Gammerdinger, W.J., Roberts, R.B., Carleton, K.L., Lien, S. and Kocher, T.D., 2019. Chromosome-scale assemblies reveal the structural evolution of African cichlid genomes. Gigascience, 8(4), p.giz030.

8. DeLorenzo L, Mathews D, Brandon AA, Joglekar M, Carmona Baez A, Moore EC, Ciccotto PJ, Roberts NB, Roberts RB, Powder KE. Genetic basis of ecologically relevant body shape variation among four genera of cichlid fishes. Mol Ecol. 2023 Jul;32(14):3975–3988.

9. Duque-Correa MJ, Codron D, Meloro C, McGrosky A, Schiffmann C, Edwards MS, Clauss M. Mammalian intestinal allometry, phylogeny, trophic level and climate. Proc Biol Sci. 2021 Feb 10;288(1944):20202888. doi: 10.1098/rspb.2020.2888.

10. Faber-Hammond JJ, Coyle KP, Bacheller SK, Roberts CG, Mellies JL, Roberts RB, Renn SCP. The intestinal environment as an evolutionary adaptation to mouthbrooding in the Astatotilapia burtoni cichlid. FEMS Microbiol Ecol. 2019 Mar 1;95(3):fiz016.

11. Feller AF, Seehausen O. Genetic architecture of adaptive radiation across two trophic levels. Proc Biol Sci. 2022 May 11;289(1974):20220377. doi: 10.1098/rspb.2022.0377. Epub 2022 May 4. PMID: 35506225; PMCID: PMC9065965.

12. Fryer, G. & T. D. Iles, 1972. The Cichlid Fishes of the Great Lakes of Africa. T.F.H. Publications, Neptune City.

13. Genner, M.J. and Turner, G.F., 2005. The mbuna cichlids of Lake Malawi: a model for rapid speciation and adaptive radiation. Fish and fisheries, 6(1), pp.1–34.

14. German, D.P. and M.H. Horn, 2006. Gut length and mass in herbivorous and carnivorous prickleback fishes (Teleostei: Stichaeidae): ontogenetic, dietary, and phylogenetic effects. Mar. Biol. 148:1123–1134.

15. Gibson, G. and Dworkin, I., 2004. Uncovering cryptic genetic variation. Nature Reviews Genetics, 5(9), pp.681–690.

16. Greene LK, McKenney EA, Gasper W, Wrampelmeier C, Hayer S, Ehmke EE, Clayton JB. Gut Site and Gut Morphology Predict Microbiome Structure and Function in Ecologically Diverse Lemurs. Microb Ecol. 2022 May 14.

17. Griffen BD, Mosblack H. Predicting diet and consumption rate differences between and within species using gut ecomorphology. J Anim Ecol. 2011 Jul;80(4):854–63.

18. JMP®, Version 17.0.0, SAS Institute Inc., Cary, NC, 1989–2023.

19. Jiang, C. and Zeng, Z.B., 1995. Multiple trait analysis of genetic mapping for quantitative trait loci. Genetics, 140(3), pp.1111–1127.

20. Kao, C.H., Zeng, Z.B. and Teasdale, R.D., 1999. Multiple interval mapping for quantitative trait loci. Genetics, 152(3), pp.1203–1216.

21. Karasov WH, Douglas AE. Comparative digestive physiology. Compr Physiol. 2013 Apr;3(2):741–83.

22. Konings, A. F., 2007. Malawi Cichlids in Their Natural Habitat. Cichlid Press, El Paso.

23. Li, H., 2013. Aligning sequence reads, clone sequences and assembly contigs with BWA-MEM. arXiv preprint arXiv:1303.3997.

24. Mackay TF. Epistasis and quantitative traits: using model organisms to study gene-gene interactions. Nat Rev Genet. 2014 Jan;15(1):22–33.

25. Malinsky, M., Svardal, H., Tyers, A.M., Miska, E.A., Genner, M.J., Turner, G.F. and Durbin, R., 2018. Whole-genome sequences of Malawi cichlids reveal multiple radiations interconnected by gene flow. Nature ecology & evolution, 2(12), pp.1940–1955.

26. Mank, J. Population genetics of sexual conflict in the genomic era. Nat Rev Genet 18, 721–730 (2017). 10.1038/nrg.2017.83

27. Moore EC, Ciccotto PJ, Peterson EN, Lamm MS, Albertson RC, Roberts RB. Polygenic sex determination produces modular sex polymorphism in an African cichlid fish. Proc Natl Acad Sci U S A. 2022 Apr 5;119(14):e2118574119.

28. Naya, D.E., Karasov, W.H., & Bozinovic, F. (2007). Phenotypic plasticity in laboratory mice and rats: a meta-analysis of current ideas on gut size flexibility. Evolutionary Ecology Research, 9, 1363–1374.

29. Nichols, P., Genner, M.J., Van Oosterhout, C., Smith, A., Parsons, P., Sungani, H., Swanstrom, J. and Joyce, D.A., 2015. Secondary contact seeds phenotypic novelty in cichlid fishes. Proceedings of the Royal Society B: Biological Sciences, 282(1798), p.20142272.

30. Olsson J, Quevedo M, Colson C, Svanbäck. 2007. Gut length plasticity in perch: into the bowels of resource polymorphisms. Biol J Linnean Soc 90:517–523.

31. O’Quin KE, Hofmann CM, Hofmann HA, Carleton KL. Parallel evolution of opsin gene expression in African cichlid fishes. Mol Biol Evol. 2010 Dec;27(12):2839–54.

32. Pfennig DW. Polyphenism in spadefoot toad tadpoles as a locally adjusted evolutionary stable strategy. Evolution. 1992 Oct;46(5):1408–1420.

33. R Core Team (2021). R: A language and environment for statistical computing. R Foundation for Statistical Computing, Vienna, Austria. https://www.R-project.org/.

34. Reinthal, P.N., 1989. The gross intestine morphology of a group of rock-dwelling Cichlid (pisces, teleostei) from Lake Malawi. Netherlands Journal of Zoology. 39(3-4): 208–225.

35. Ribbink, A. J., B. A. Marsh, A. C. Marsh, A. C. Ribbink & B. J. Sharp, 1983. A preliminary survey of the cichlid fishes of rocky habitats in Lake Malawi. South African Journal of Zoology 3: 149–310.

36. Riddle MR, Aspiras A, Damen F, McGaugh S, Tabin JA, Tabin CJ. Genetic mapping of metabolic traits in the blind Mexican cavefish reveals sex-dependent quantitative trait loci associated with cave adaptation. BMC Ecol Evol. 2021 May 21;21(1):94.

37. Roberts, R.B., Ser, J.R. and Kocher, T.D., 2009. Sexual conflict resolved by invasion of a novel sex determiner in Lake Malawi cichlid fishes. Science, 326(5955), pp.998–1001.

38. Roberts RB, Hu Y, Albertson RC, Kocher TD. Craniofacial divergence and ongoing adaptation via the hedgehog pathway. Proc Natl Acad Sci U S A. 2011 Aug 9;108(32):13194–9.

39. Ronco, F., Roesti, M. and Salzburger, W., 2019. A functional trade-off between trophic adaptation and parental care predicts sexual dimorphism in cichlid fish. Proceedings of the Royal Society B, 286(1909), p.20191050.

40. Ser JR, Roberts RB, Kocher TD. Multiple interacting loci control sex determination in lake Malawi cichlid fish. Evolution. 2010 Feb 1;64(2):486–501. doi: 10.1111/j.1558-5646.2009.00871.x. Epub 2009 Oct 23. PMID: 19863587; PMCID: PMC3176681.

41. Silva, L.D.C.E., Wang, S. and Zeng, Z.B., 2012. Composite interval mapping and multiple interval mapping: procedures and guidelines for using Windows QTL Cartographer. In Quantitative trait loci (QTL) (pp. 75-119). Humana Press.

42. Starck JM. 2005. Structural flexibility of the digestive system of tetrapods: patterns and processes at the cellular and tissue level. In: Starck JM, Wang T, editors. Physiological and ecological adaptations to feeding in vertebrates. Enfield: Science Publishers Inc. p 175–200.

43. Waddington, C.H., 1942. Canalization of development and the inheritance of acquired characters. Nature, 150(3811), pp.563–565.

44. Wagner, C.E., McIntyre, P.B., Buels, K.S., Gilbert, D.M. and Michel, E., 2009. Diet predicts intestine length in Lake Tanganyika’s cichlid fishes. Functional Ecology, 23(6), pp.1122–1131.

45. Wang S., C. J. Basten, and Z.-B. Zeng (2012). Windows QTL Cartographer 2.5. Department of Statistics, North Carolina State University, Raleigh, NC. http://statgen.ncsu.edu/qtlcart/WQTLCart.htm

46. Wickham, H., Averick, M., Bryan, J., Chang, W., McGowan, L.D.A., François, R., Grolemund, G., Hayes, A., Henry, L., Hester, J. and Kuhn, M., 2019. Welcome to the Tidyverse. Journal of open source software, 4(43), p.1686.

47. Yawitz TA, Barts N, Kohl KD. Comparative digestive morphology and physiology of five species of Peromyscus under controlled environment and diet. Comp Biochem Physiol A Mol Integr Physiol. 2022 Sep;271:111265.

48. Zeng, Z.B., 1994. Precision mapping of quantitative trait loci. Genetics, 136(4), pp.1457–1468.

